# TIGIT is upregulated by HIV-1 infection and marks a highly functional adaptive and mature subset of natural killer cells

**DOI:** 10.1101/764217

**Authors:** Elena Vendrame, Christof Seiler, Thanmayi Ranganath, Nancy Q. Zhao, Rosemary Vergara, Michel Alary, Annie-Claude Labbé, Fernand Guédou, Johanne Poudrier, Susan Holmes, Michel Roger, Catherine A. Blish

## Abstract

**Objective:** Our objective was to investigate the mechanisms that govern natural killer (NK) cell responses to HIV, with a focus on specific receptor-ligand interactions involved in HIV recognition by NK cells.

**Design and Methods:** We first performed a mass cytometry-based screen of NK cell receptor expression patterns in healthy controls and HIV^+^ individuals. We then focused mechanistic studies on the expression and function of T cell immunoreceptor with Ig and ITIM domains (TIGIT).

**Results:** The mass cytometry screen revealed that TIGIT is upregulated on NK cells of untreated HIV^+^ women, but not in antiretroviral-treated women. TIGIT is an inhibitory receptor that is thought to mark exhausted NK cells; however, blocking TIGIT did not improve anti-HIV NK cell responses. In fact, the TIGIT ligands CD112 and CD155 were not upregulated on CD4^+^ T cells *in vitro* or *in vivo*, providing an explanation for the lack of benefit from TIGIT blockade. TIGIT expression marked a unique subset of NK cells that express significantly higher levels of NK cell activating receptors (DNAM-1, NTB-A, 2B4, CD2) and exhibit a mature/adaptive phenotype (CD57^hi^, NKG2C^hi^, LILRB1^hi^, FcRγ^lo^, Syk^lo^). Furthermore, TIGIT^+^ NK cells had increased responses to mock-infected and HIV-infected autologous CD4^+^ T cells, and to PMA/ionomycin, cytokine stimulation and the K562 cancer cell line.

**Conclusions:** TIGIT expression is increased on NK cells from untreated HIV^+^ individuals. Although TIGIT does not participate directly in NK cell recognition of HIV, it marks a population of mature/adaptive NK cells with increased functional responses.

## INTRODUCTION

Natural killer (NK) cells are among the first responders to viral infections and can swiftly recognize and kill virus-infected cells [1]. These responses are traditionally thought to be nonspecific, as NK cell function is primarily mediated by the expression of germ-line encoded receptors rather than antigen-specific receptors [2]. However, recent evidence has revealed that NK cells are capable of antigen-specific adaptive responses against viruses, such as cytomegalovirus, Epstein-Barr Virus, Varicella Zoster Virus, and influenza virus [1,3–9].

HIV infection profoundly alters the NK cell compartment, with expansion of a CD56^-^CD16^+^ subpopulation [10,11], downregulation of several activating NK cell receptors, including FcRγ (FcεRIγ), NKp30, and NKp46 [12–14], and impairment of NK cell function [15–18]. NK cells also play a critical role in the immune response against HIV [19,20]. NK cells may protect from HIV acquisition, as increased NK cell activity is associated with lower risk of acquiring HIV in highly exposed individuals [21–23]. Consistent with this, genotypes of specific NK cell receptors and human leukocyte antigens are enriched in exposed seronegative individuals [21,24–26]. The expression of specific NK cell receptor/HLA class I ligand pairs (KIR3DL1/KIR3DS1 with HLABw4-80I) is also associated with slower disease progression [27,28] and NK cells from long term non-progressors show increased function compared to HIV^+^ typical progressors [12,29]. Altogether, these data raise the possibility that specific NK receptor-ligand interactions may contribute to HIV control and could be used as targets to improve HIV-specific NK cell function.

T cell immunoreceptor with Ig and ITIM domains (TIGIT) is an inhibitory receptor expressed on T cells and NK cells [30,31]. The TIGIT inhibitory signal is mediated by ligation with its high affinity ligand, the poliovirus receptor (CD155 or PVR), and its low affinity ligand CD112 (Nectin-2 or PVRL2) [30,32,33]. TIGIT has been previously associated with CD4^+^ T cell, CD8^+^ T cell and NK cell exhaustion both in the setting of chronic viral infections and malignancy [31,34–40]. Blockade of the TIGIT/CD155/CD112 pathway to improve the function of T cells and NK cells against solid cancers is currently under investigation [41,42].

In HIV-1 infection, TIGIT marks exhausted T cells, correlates with disease progression and is decreased on CD4^+^ T cells from elite controllers [43,44]. Early initiation of antiretroviral treatment (ART) in HIV infected individuals does not return TIGIT/CD115 to normal levels on CD8^+^ T cells [37]. Additionally, the co-expression of TIGIT with immune checkpoint inhibitor PD-1 marks CD4^+^ T cells harboring latent virus [45–47]. These data suggest that the TIGIT/CD155/CD112 pathway in T cells could contribute to HIV pathogenesis.

In NK cells, less is known regarding the role of TIGIT during HIV infection. A recent report showed that TIGIT is increased on NK cells from HIV-1 infected patients and that TIGIT blockade improves NK cell responses to cytokines *ex-vivo* in these patients [48]. Here, we used blood samples from Beninese women to study the effect of HIV infection on the NK cell compartment, with a particular focus on TIGIT expression and function.

## METHODS

### Study subjects and sample processing

Cryopreserved peripheral blood mononuclear cells (PBMCs) were obtained from a study of commercial sex workers in Cotonou, Benin. HIV-1-infected women were enrolled from a sex-worker clinic and HIV-1-uninfected women were enrolled from a general health clinic, as described previously [49,50]. PBMCs were obtained from 20 untreated HIV-infected women, 20 HIV-infected women receiving ART and 10 healthy women (Table S1). Written informed consent was obtained from all subjects. The study was approved by the Comité National Provisoire d’Éthique de la Recherche en Santé in Cotonou and the Centre Hospitalier de l’Université de Montréal (CHUM) Research Ethics Committees.

For mechanistic *in vitro* studies using healthy donors, leukoreduction system (LRS) chambers were purchased from the Stanford Blood Bank. PBMCs were purified using Ficoll density gradient centrifugation and cryopreserved in fetal bovine serum (FBS) with 10% DMSO.

#### Cell isolation

For profiling PBMCs from Beninese women, 1×10^6^ PBMCs were stained for mass cytometry with the ligand antibody panel (Table S2). NK cells were purified from PBMCs by magnetic-activated isolation via negative selection (Miltenyi) and stained with the NK cell antibody panel (Table S3).

For the *in vitro* co-cultures, cryopreserved PBMCs from healthy donors were used to purify CD4^+^ T cells and NK cells by magnetic-activated isolation via negative selection (Miltenyi).

#### Antibody conjugation, mass cytometry staining and data acquisition

Antibodies were conjugated using MaxPar^®^ ×8 labeling kits (Fluidigm). To ensure antibody stability over time, antibody panels were lyophilized into single-use pellets prior to use (Biolyph). Cells were stained for mass cytometry as described previously [51,52] and resuspended in 1× EQ Beads (Fluidigm) before acquisition on a Helios mass cytometer (Fluidigm).

#### *In vitro* HIV infection

Replication competent (Q23-FL) HIV-1 was made by transfection and titrated as described [51]. CD4^+^ T cells were activated for 2 days with plate-bound anti-CD3 (clone OKT3, eBioscience, 10 μg/ml), soluble anti-CD28/CD49d (clones L293/L25, BD Biosciences, 1 μg/ml each) and phytohemagglutinin (eBioscience, 2.5 μg/ml). CD4^+^ T cells were infected overnight with Q23-FL via ViroMag R/L magnetofection (Oz Biosciences) at a multiplicity of infection of 20. HIV infection was as measured by intracellular p24.

#### TIGIT blockade assay

Cells were cultured in RPMI media containing 10% FBS (Thermo Fisher Scientific) and 1% penicillin/streptomycin/amphotericin (Thermo Fisher Scientific) (RP10). NK cells were incubated in 24-well plates at a concentration of 2×10^6^ cells/ml in for 3 days at 37°C with 5% CO2, with addition of rhIL-2 (R&D Systems, 300 IU/ml). After 3 days, NK cells were washed with fresh media and plated in 96-well plates (80,000-100,000 cells/well). Blocking mIgG1κ anti-hTIGIT antibody (eBioscience, clone MBSA43, 10 μg/ml) or a mIgG1κ isotype control (eBioscience, clone P3.6.2.8.1, 10 μg/ml) were added. Cells were incubated at 4°C for 20 minutes, washed with FACS wash buffer (PBS + 2% FBS) and resuspended in media.

#### CD112 and CD155 mRNA expression levels

RNA from resting, mock-infected or HIV-infected CD4^+^ T cells was isolated using the RNeasy Mini Kit with QIAshredder columns (Qiagen). cDNA was produced using the SuperScript III First Strand Synthesis Kit (Thermo Fisher Scientific). Quantitative PCR (qPCR) was achieved with Taqman Universal Master Mix (Applied Biosystems) with the following commercially bought Taqman Assays, using fluorescein amidite-minor groove binder (FAM-MGB) probes: Nectin-2/CD112 (Thermo Fisher Scientific), PVR/CD155 (Thermo Fisher Scientific), and Human GAPDH endogenous control (Applied Biosystems). qPCR was run on the Applied Biosystems Step One Plus Real Time PCR System. Transcription levels were normalized to GAPDH. mRNA fold change expression was calculated using the double delta Ct analysis method with resting CD4^+^ T cells as the control group [53].

#### NK cell:CD4^+^ T cell co-cultures, cytokine stimulation and determination of NK cell function

All cells were cultured in RP10. NK cells (80,000-100,000/well) were incubated with HIV-infected or mock-infected autologous CD4^+^ T cells (320,000/well at a 1:4 effector:target ratio), 0.4 μl/200μl of cell stimulation cocktail (phorbol 12-myristate 13-acetate (PMA) and ionomycin, eBioscience), a cocktail of IL-12, IL-15 and IL-18 (rhIL-12, R&D Systems, 5ng/ml; rhIL-15, Pepro-Tech, 20ng/ml; rhIL-18, R&D Systems, 0.5 μg/ml), or with the cancer cell line K562 (ATCC, used at passage 2-10, 300,000 cells/well at a 1:3 effector:target ratio). Cells were incubated for 4 hours at 37°C with 5% CO2, in the presence of brefeldin, monensin (eBioscience) and anti-CD107a-APC-H7 antibody (BD Biosciences, Clone H4A3). After incubation, cells were stained with LIVE/DEAD^™^ Fixable Yellow Staining Kit (Thermo Fisher Scientific) for 20 minutes at room temperature, then fixed (BD FACS Lyse), permeabilized (BD FACS Perm II) and stained with anti-IFN-γ-V450 (BD Biosciences, Clone B27), anti-TIGIT-APC (eBioscience, Clone MBSA43), anti-CD3-PerCP-Cy5.5 (Biolegend, Clone UCHT1), anti-HIVp24-FITC (Beckman Coulter, Clone KC57) for 30 minutes at 4°C. Data was acquired on a MACSQuant^®^ Analyzer Flow Cytometer (Miltenyi).

#### Data analysis

Bead normalization was performed before downstream analyses (https://github.com/nolanlab/bead-normalization) [54]. To ascertain the quality of the stains, data were visualized using FlowJo v10.5.3. FAS-L, Ki-67 and CXCR6 were excluded from subsequent analyses due to poor staining. The original data were uploaded in ImmPort (www.immport.org). Serial negative gating was performed to identify NK cells from any contaminating cells remaining after purification (Figure S1a). Examples of stains for NK markers are shown in Figure S2a. Samples with cell number <1000 were excluded from analyses; the number of subjects used for each analysis is specified in the figure legend. The open source statistical package R (https://www.r-project.org/) was used for all statistical analyses [55]. Normalized signal intensities were transformed using the asinh function with a cofactor of 5 prior to generalized linear model (GLM) analysis, multidimensional scaling (MDS) and Uniform Manifold Approximation and Projection (UMAP) visualization. For the GLM analysis, we used the custom-made package “CytoGLMM” [56,57]. This package implements a generalized linear model with bootstrap resampling to identify markers predictive of clinical treatment groups. The “CytoGLMM” package was also used to create a MDS projection. For comparison between groups, we used the Wilcoxon rank-sum test. When appropriate, p-values were adjusted for multiple comparisons using the Benjamini-Hochberg method. UMAP embeddings were calculated using the R package “uwot”, with n_neighbor = 5 and min_dist = 0.2, as described [58]. Gated NK cells from all samples were pooled into treatment groups, to facilitate visualization. Embeddings were visualized in Cytobank (www.cytobank.org).

### RESULTS

#### TIGIT is upregulated on NK cells from untreated HIV-infected women

To investigate the effect of HIV-1 infection on NK cells, we used CyTOF to profile the NK cells of twenty untreated HIV-1-infected women, twenty HIV-1 infected women on ART and ten healthy women from the same geographical area (Figure 1a). To analyze this multidimensional dataset, we first used a custom-made generalized linear model with bootstrap resampling (Figure 1b). This model confirmed that HIV-1 infection is associated with a profound alteration of the NK cell phenotype, with higher levels of CD38 and lower levels of Siglec-7 predicting chronic HIV-1 infection as previously described [59,60]. This analysis also confirmed a recent finding that checkpoint inhibitor TIGIT is upregulated in untreated HIV-1-infected individuals [48]. Additionally, our data revealed that TIGIT is not elevated on NK cells from antiretroviral-treated HIV-1-infected women (Figure 1b and 1c).

**Figure 1:**
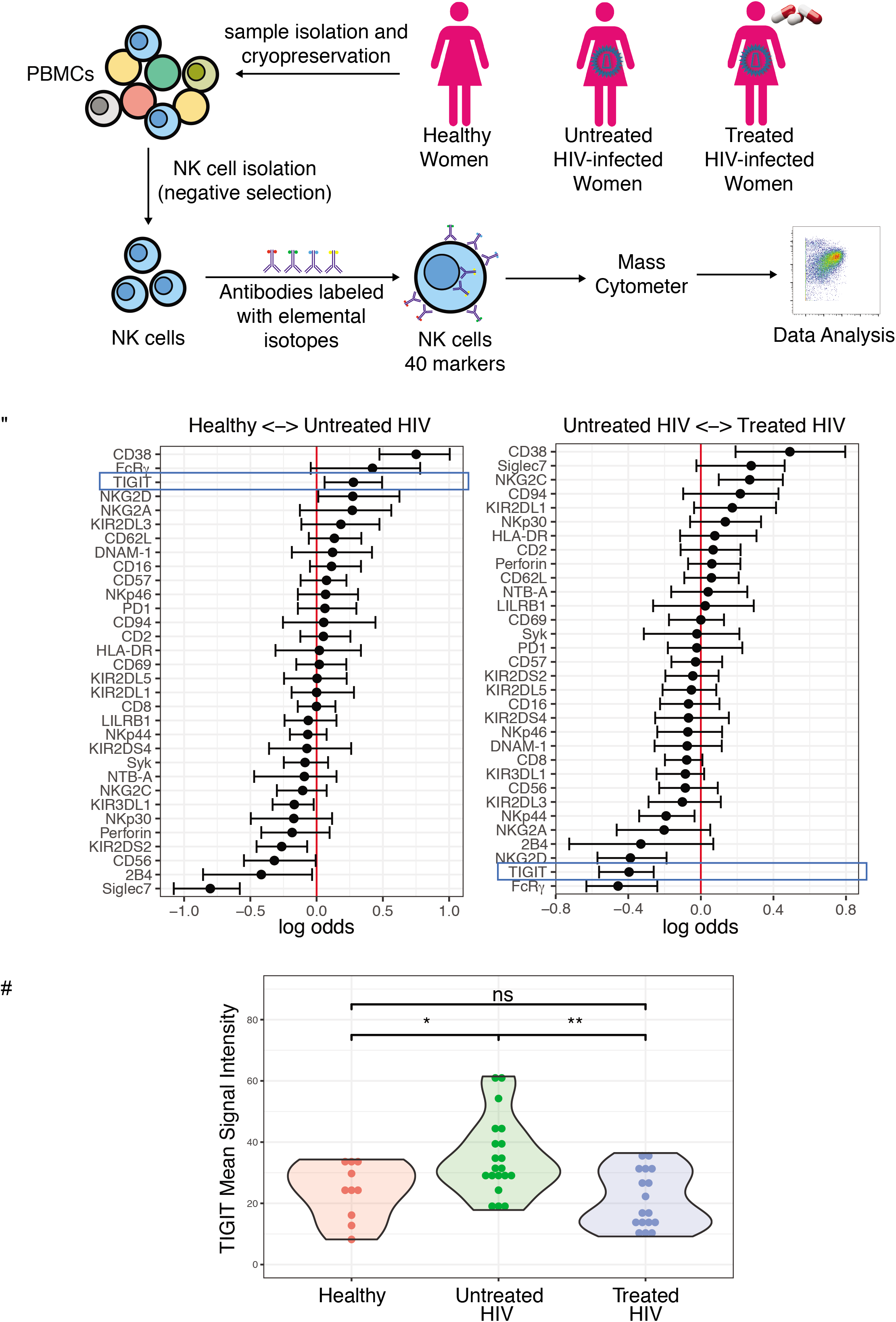
TIGIT is upregulated on NK cells from untreated HIV-infected women. **(A)** Schematic of study design and experiment. **(B)** A generalized linear model with bootstrap resampling was used to identify markers predictive of the study groups. Log-odds are logarithm of ratios of the probability that a cell belongs to each study subject. An increase in the parameter coefficient corresponds to the strength of the classification power, with the 95% confidence interval represented by line surrounding the point estimate. **(C)** Mean Signal Intensity of TIGIT on NK cells from healthy women (n=10), untreated HIV^+^ women (n=20), treated HIV^+^ women (n=17). * = unadjusted p-value≤0.05, ** = unadjusted p-value≤0.01, ns = non-significant.

#### TIGIT expression is increased in CD56^dim^ and CD56^-^ NK cells from untreated HIV-1^+^ women

To further explore the nature of this increased TIGIT expression during HIV-infection, we evaluated TIGIT expression within different NK cell subpopulations. NK cells are classically divided into three distinct subpopulations: CD56^bright^, which secrete cytokines at high levels, CD56^dim^, a more cytotoxic subpopulation, and CD56^-^, a subpopulation of NK cells that are thought to be functionally impaired and are expanded during chronic viral infections, including HIV [10,61–63]. Here we confirmed that CD56^-^ NK cells are increased in HIV-infected individuals, regardless of treatment status (Figure 2a and 2b). In addition, we found that TIGIT expression is higher on CD56^-^ NK cells from untreated HIV-1 infected women compared to healthy controls, and that TIGIT expression is increased in the CD56^dim^ subpopulation in untreated HIV-1^+^ women compared to treated women (Figure 2c).

**Figure 2.**
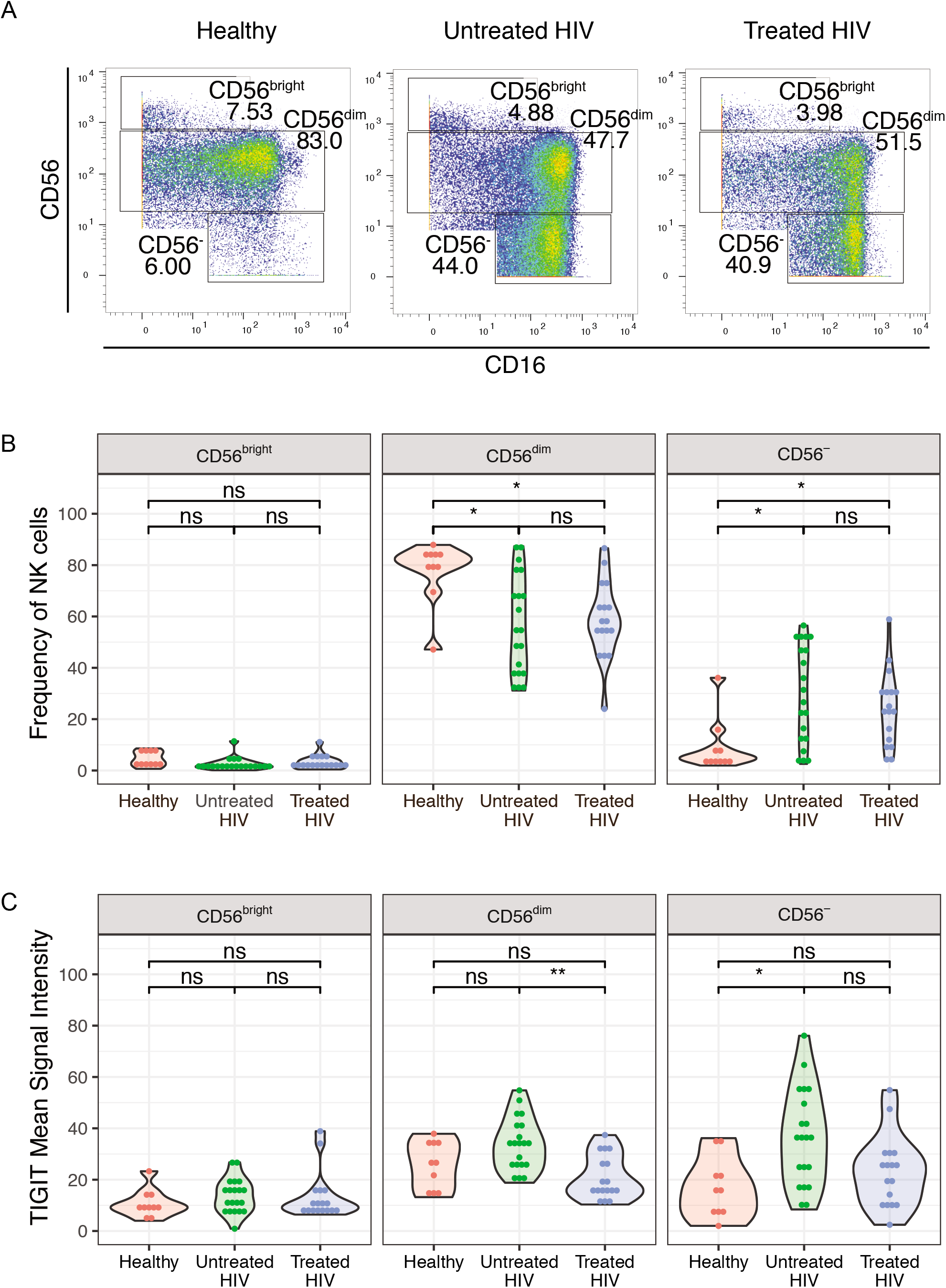
TIGIT expression is increased in CD56^dim^ and CD56^-^ NK cells of untreated HIV-1^+^ women. **(A)** Representative CyTOF plots of gating of NK cells based on CD56 expression. (B) Frequency of CD56^bright^, CD56^dim^ and CD56^-^ NK cells in healthy women (n=10), untreated HIV^+^ women (n=20), treated HIV^+^ women n=17). (C) Mean Signal Intensity of TIGIT on NK cell subsets from healthy women (n=10), untreated HIV^+^ women (n=20), treated HIV^+^ women (n=17). * = adjusted p-value≤0.05, ** = adjusted p-value≤0.01, ns = non-significant.

#### TIGIT blockade does not alter HIV-specific NK cell responses

TIGIT blockade increases *ex vivo* NK cell responses to cytokines in HIV-1 infected women [48] and elicits improved anti-tumor responses in mouse models [35]. To explore the effect of TIGIT blockade on NK cell responses against HIV *in vitro*, we treated NK cells from healthy donors with an anti-TIGIT monoclonal antibody and measured NK cell responses (CD107a and IFN-γ) by flow cytometry after incubation with mock-infected or HIV-infected autologous CD4^+^ T cells (Figure 3a). Blockade of TIGIT did not improve NK cell responses to mock-infected or HIV-infected CD4^+^ T cells, when compared to untreated NK cells or cells treated with an isotype control (Figures 3b and 3c).

**Figure 3.**
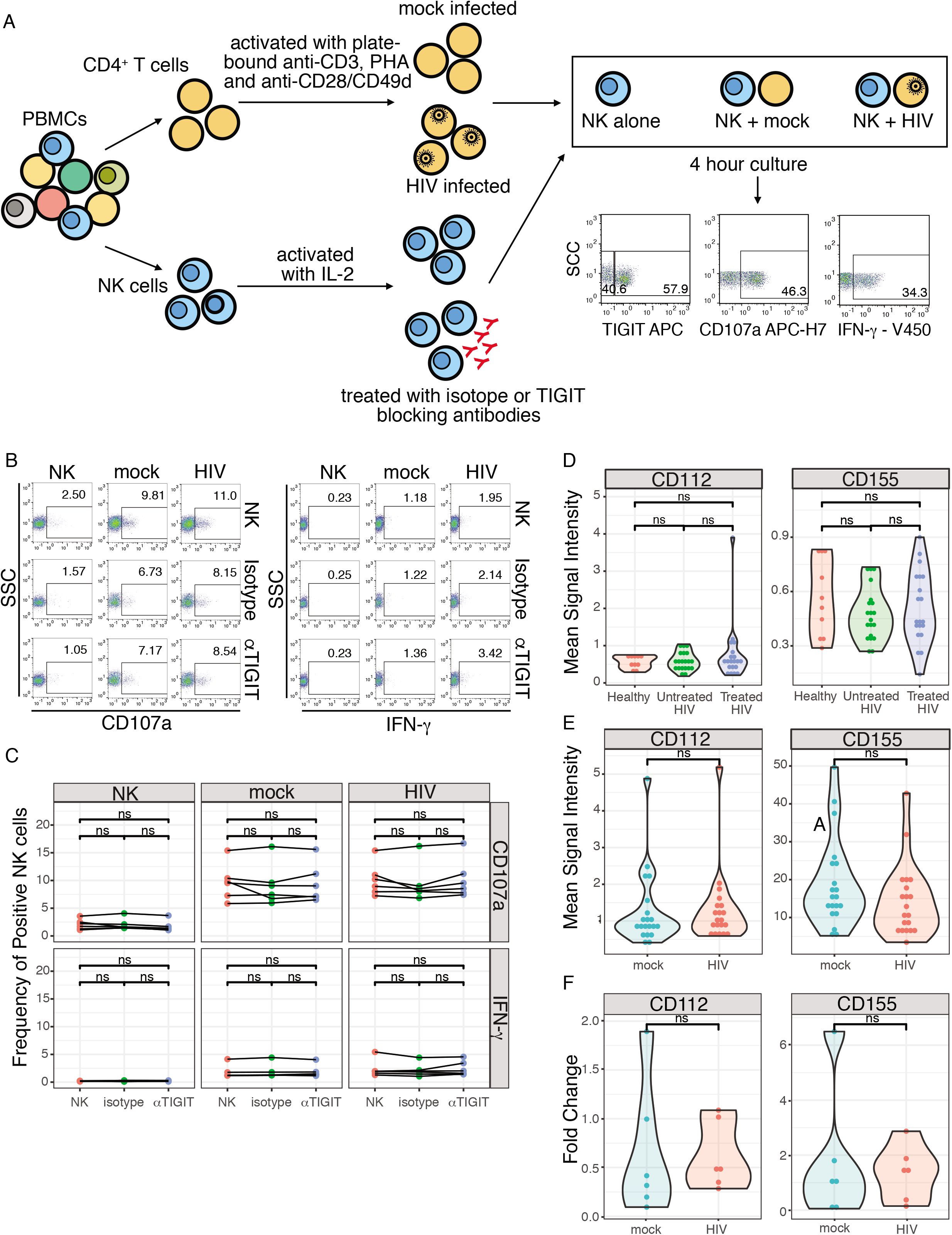
TIGIT blockade does not alter HIV-specific NK cell responses to HIV and TIGIT ligands are not upregulated by during HIV infection. **(A)** Schematic of TIGIT blockade assay and NK cell:CD4^+^ T cell co-cultures. **(B)** Representative flow cytometry plots of CD107a and IFN-γ production, expressed as frequency of positive cells, by NK cells alone (NK) or after coculture with mock-infected (mock) or HIV-infected (HIV) autologous CD4^+^ T cells. Cells were incubated alone (NK), pre-treated with isotype control (Isotype) or a blocking anti-TIGIT antibody (αTIGIT). **(C)** Summary plot of frequency of CD107a^+^ and IFN-γ^+^ NK cells after 4 hour co-culture with mock-infected or HIV-infected autologous CD4^+^ T cells. (D) Mean Signal Intensity of CD112 and CD155 expression measured by mass cytometry in healthy women (n=9), untreated HIV^+^ women (n=20), treated HIV^+^ women (n=17). (E) Mean Signal Intensity of CD112 and CD155 expression measured by mass cytometry in mock-infected and HIV-infected CD4^+^ T cells. (F) Fold change in the *NECTIN2* (CD112) and *PVR* (CD155) transcript levels, normalized to *GAPDH*, in mock-infected and HIV-infected CD4^+^ T cells, relative to resting CD4^+^ T cells. ns = non-sigificant.

#### TIGIT ligands are not upregulated by HIV infection

The TIGIT inhibitory signal is mediated by ligation with CD155 (high affinity) or CD112 (low affinity) [30,32]. To investigate the role of the TIGIT-CD155 and TIGIT-CD112 pathways in the setting of HIV infection, we measured the expression of CD155 and CD112 on HIV-infected CD4^+^ T cells *ex vivo* and *in vitro.* CD155 and CD112 were both expressed at low levels on CD4^+^ T cells from HIV^+^ women and there was no difference in expression between healthy women, untreated HIV^+^ women and treated HIV^+^ women (Figures 3d and S2b). CD155 and CD112 were not differentially expressed in HIV-infected or mock-infected CD4^+^ T cells *in vitro* when measured by CyTOF (Figures 3e and S2c). Additionally, CD112 and CD115 transcription levels did not differ between mock-infected and HIV-infected CD4+ T cells (Figure 3f) with a median fold change of HIV relative to mock-infected CD4^+^ T cells 1.15 (interquartile range = 0.94-2.4) for CD155 and 1.07 (interquartile range = 0.93-3.13) for CD112.

#### TIGIT expression is increased in adaptive/mature NK cells

To characterize NK cells with increased TIGIT expression, we gated on TIGIT and compared TIGIT^+^ and TIGIT^-^ NK cells from our Beninese women (Figure 4a). Using MDS, we found that TIGIT expression marks a distinct population of NK cells that was not explained by HIV infection or treatment status (Figure 4b-d). We then focused our analysis on features distinguishing

**Figure 4.**
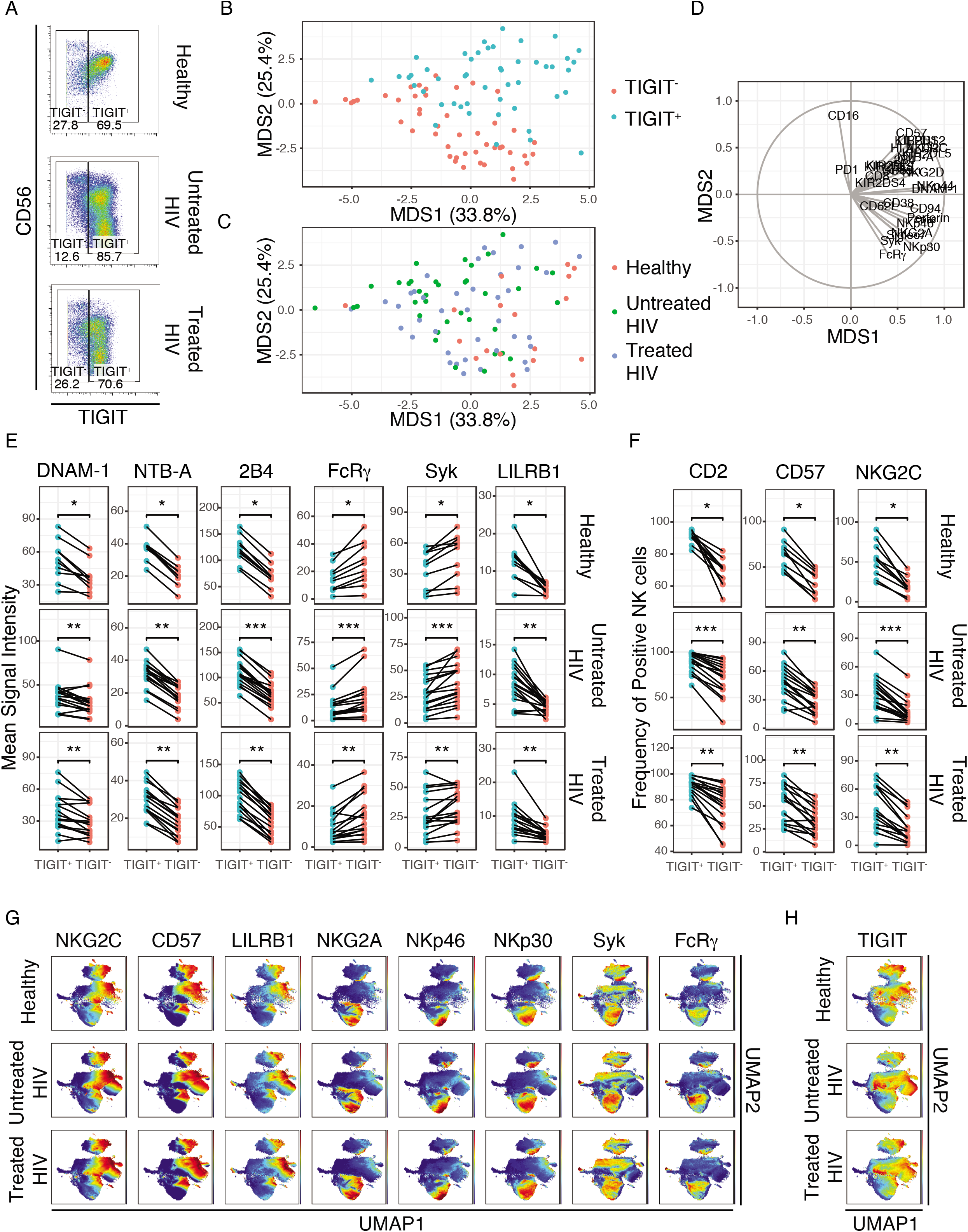
TIGIT expression is increased in adaptive/mature NK cells. **(A)** Representative CyTOF plots of gating for TIGIT^+^ and TIGIT^-^ NK cells from healthy women (n=9), untreated HIV^+^ women (n=17), treated HIV^+^ women (n=16). **(B)** Multidimensional Scaling (MDS) of TIGIT^+^ and TIGIT^-^ NK cells colored by TIGIT expression. **(C)** MDS of TIGIT^+^ and TIGIT^-^ NK cells colored by group (Healthy, Untreated HIV and Treated HIV). **(D)** Vectors driving the variance of the MDS. **(E)** Mean Signal Intensity of DNAM-1, NTB-A, 2B4, FcRγ, Syk and LILRB1 expression in TIGIT^+^ and TIGIT^-^ NK cells from healthy women, untreated HIV^+^ women and treated HIV^+^ women. **(F)** Frequency of CD2^+^, CD57^+^ and NKG2C^+^ NK cells from TIGIT^+^ and TIGIT^-^ NK cells from healthy women, untreated HIV^+^ women and treated HIV^+^ women. **(G)** UMAP plot of pooled NK cells from healthy women, untreated HIV^+^ women and treated HIV^+^ women colored by markers of adaptive/mature NK cells (NKG2C, CD57, LILRB1, NKG2A, NKp46, NKp30, Syk, FcRγ). **(H)** UMAP plot of pooled NK cells from healthy women, untreated HIV^+^ women and treated HIV^+^ women, colored by TIGIT expression. * = adjusted p-value≤0.05, ** = adjusted p-value≤0.01, *** = adjusted p-value≤0.001.

TIGIT^+^ and TIGIT^-^ NK cells. We found that TIGIT^+^ NK cells express higher levels of activating markers (DNAM-1, NTB-A, 2B4, CD2) and exhibit an adaptive/mature phenotype (CD57^hi^, NKG2C^hi^, LILRB1 (ILT2, CD85j)^hi^, FcRγ^lo^, Syk^lo^) (Figures 4e and 4f). To follow-up on this finding, we used UMAP to visualize NK cell subsets and found that adaptive/mature NK cells (CD57^hi^, NKG2C^hi^, LILRB1^hi^, NKG2A^lo^, NKp46^lo^, NKp30^lo^, FcRγ^lo^, Syk^lo^) were separated on the right side of the UMAP1 axis (Figure 4g). Additionally, when we colored the UMAP plot by TIGIT expression, we found that TIGIT expression co-localized with markers of adaptive/mature NK cells (Figure 4h).

#### TIGIT expression is associated with increased NK cell responses

As mature NK cells are typically associated with higher functional activity, we sought to better characterize the function of TIGIT^+^ and TIGIT^-^ NK cells. We isolated NK cells from healthy donors and cultured them with mock-infected or HIV infected autologous CD4^+^ T cells (Figure 5a). We then measured the functional responses (CD107a and IFN-γ) of TIGIT^+^ and TIGIT^-^ NK cells by flow cytometry (Figures 5b-e). We found that TIGIT^+^ NK cells had increased responses to mock-infected and HIV-infected autologous CD4^+^ T cells (Figures 5b and 5c). To better define whether this difference was specific to HIV, we also measured the responses to PMA/ionomycin, a cocktail of IL-12, IL-15 and IL-18, and the leukemia cell line K562 (Figures 5a, 5d and 5e). TIGIT^+^ NK cells also exhibited increased responses to PMA/ionomycin, cytokine and tumor cells, compared to TIGIT^-^ NK cells (Figures 5d and 5e).

**Figure 5.**
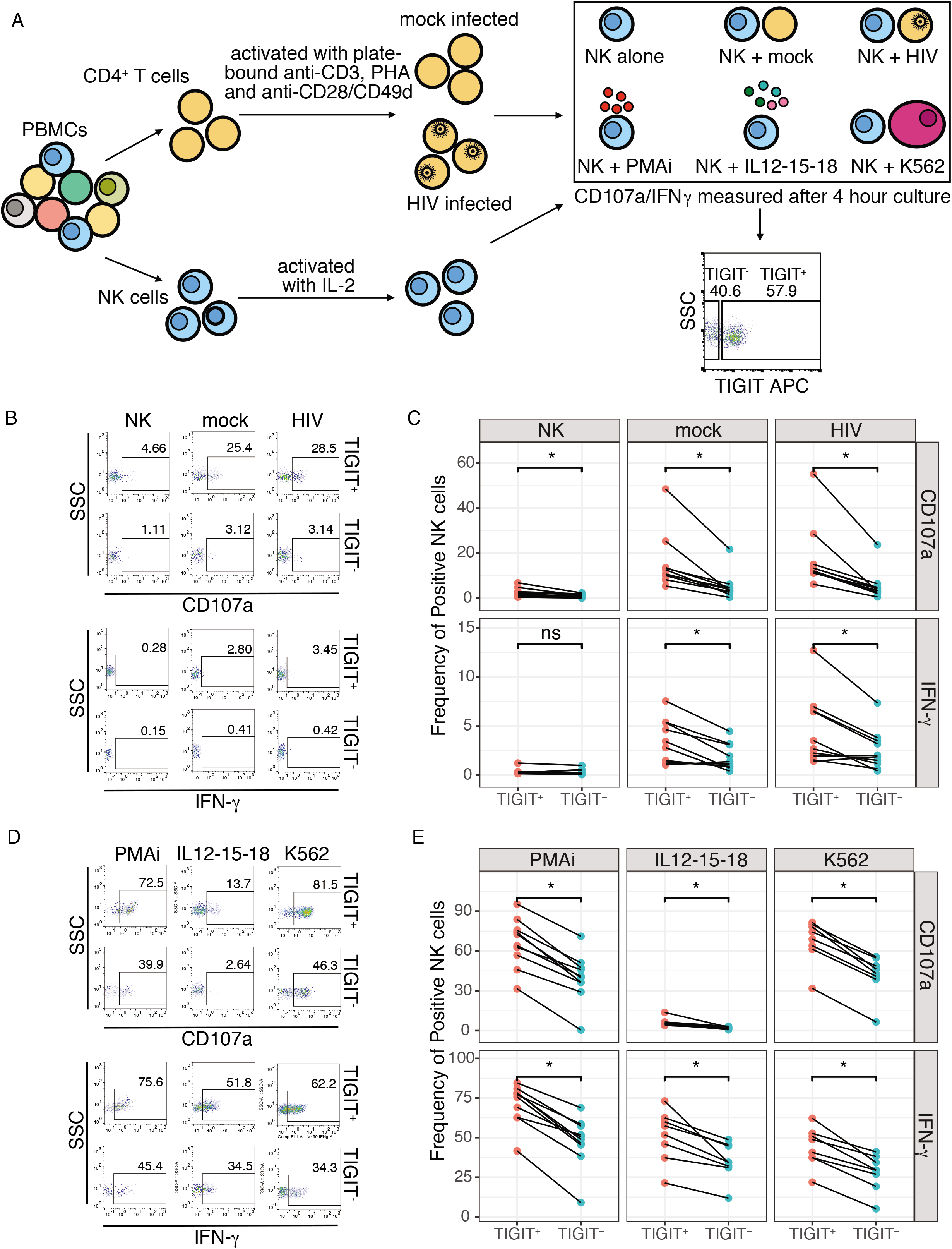
TIGIT expression is associated with increased NK cell responses. **(A)** Schematic of CD4^+^ T cell:NK cell co-cultures or stimulation and representative flow cytometry plot of gating of TIGIT^+^ and TIGIT^-^ NK cells. **(B)** Representative flow cytometry plots of CD107a and IFN-γ production, expressed as frequency of positive cells, by TIGIT^+^ and TIGIT^-^ NK cells after 4 hour co-culture with mock-infected (mock) or HIV-infected (HIV) autologous CD4^+^ T cells. **(C)** Summary plot of the frequency of TIGIT-gated CD107a^+^ and IFN-γ^+^ NK cells after 4 hour coculture with mock-infected or HIV-infected autologous CD4^+^ T cells. **(D)** Representative flow cytometry plots of CD107a and IFN-γ production by TIGIT^+^ and TIGIT^-^ NK cells after 4 hour stimulation with PMA/ionomycin (PMA/i), a cocktail of IL12, IL-15 and IL-18 (IL12-15-18) or the leukemia cell line K562 (K562). (E) Summary plot of the frequency of TIGIT-gated CD107a^+^ and IFN-γ^+^ NK cells after 4 hour stimulation with PMA/ionomycin (PMA/i), a cocktail of IL12, IL-15 and IL-18 (IL12-15-18) or the leukemia cell line K562 (K562). * = adjusted p-value≤0.05, ns = non-sigificant.

### DISCUSSION

NK cells undergo significant changes during chronic HIV infection [16,64], with expansion of a hypofunctional CD56^-^CD16^+^ NK cell subpopulation [10,11], downregulation of several activating NK cell receptors [12–14], and adaptive reconfiguration [65]. Some of these changes are not observed after treatment or in the setting of natural HIV control [66], indicating that diminished NK cell function could contribute to disease pathogenesis. To better understand the role of different NK cell receptors in the pathogenesis of HIV, we used CyTOF, a technique that allows the detection of over 40 receptors simultaneously, to profile the NK cells from infected and uninfected HIV^+^ women. A particularly striking finding was that TIGIT is upregulated in untreated HIV infection, yet its expression is normal in the setting of ART. TIGIT upregulation on NK cells from HIV-infected male patients has been recently described [48]. Here, we demonstrate that TIGIT is upregulated in HIV-infected women and that this upregulation does not occur in women receiving ART. Further, TIGIT upregulation is more prominent on CD56^-^/CD16^+^ NK cells, a subpopulation thought to be functionally exhausted [10,11].

TIGIT is an inhibitory receptor that has been associated with hypofunctional/exhausted NK cells in cancer patients and mouse models of solid tumors [31,35] and with exhausted CD8^+^ T cells in hematologic malignancy and in HIV infection [36,39,43]. As a result, several studies have explored the use of TIGIT blockade to rescue T and NK cell function. Blocking TIGIT is associated with improved anti-tumoral response in tumor-bearing mice and in patients with colorectal cancer [35], and with improved NK cell responses to cytokine stimulation in HIV^+^ patients [48]. However, we found that TIGIT blockade does not improve HIV-specific antiviral function in healthy donors. This discrepancy could be attributed by the lower expression of TIGIT by healthy individuals, as high expression levels are likely required to see the results of blockade.

The lack of a benefit for TIGIT blockade may also be explained by the fact that TIGIT is simply not engaged during NK cell recognition of HIV-infected cells. We found that TIGIT ligands CD155 and CD112 are not upregulated on CD4^+^ T cells by HIV infection in *vitro*, consistent with previous reports [67–69]. We also found that TIGIT ligands were not upregulated in HIV^+^ women, though this differs from the findings of Yin et al. [48], who reported elevated CD155 expression on CD4^+^ T cells from HIV-infected individuals. Notably, Yin et al. found that CD155 levels were very low and that blocking CD155 did not improve NK cell responses. Thus, together the data suggest that the TIGIT/CD155/CD112 pathway may not be directly involved in NK cell responses to HIV-infected cells.

While we did not find that TIGIT itself plays a key role in immune response to HIV, we noted that TIGIT marks a distinct NK subpopulation and is coexpressed with several activation markers (DNAM-1, NTB-A, 2B4, CD2). We also show that in our cohort of HIV-infected women, TIGIT^+^ NK cells exhibit a more adaptive/mature phenotype (CD57^hi^, NKG2C^hi^, LILRB1^hi^, FcRγ^lo^, Syk^lo^ and adaptive NK cells demonstrated higher TIGIT expression in unsupervised clustering via UMAP. This is consistent with the notion that NK cells undergo an adaptive reconfiguration during HIV infection [65]. Notably, Sarhan et al. demonstrated that TIGIT was expressed at lower levels in adaptive NK cells from healthy donors [70]. Our finding that TIGIT is expressed more highly on adaptive NK cells could reflect unique differentiation pathways driven by HIV infection. Alternatively, the differences in TIGIT expression between our studies may be due to differential downregulation of TIGIT in response to cytokine stimulation, given that Sarhan et al. evaluated cytokine-treated NK cells rather than PBMCs directly *ex vivo.*

Adaptive NK cells show increased responses to FcγRIIIa triggering [71–73], are expanded in HIV-infected patients and robustly respond to HIV peptides [14]. Here we demonstrate that TIGIT^+^ NK cells have higher functional activity against mock-infected and HIV-infected CD4^+^ T cells, compared to TIGIT^-^ NK cells. Further, higher functional responses also occur in the presence of non-HIV stimuli (PMA/ionomycin, cytokines and K562). This increased activity could be explained by the co-expression of activating receptors, which may overwhelm the inhibitory signal provided by TIGIT, or by changes in the activation threshold during adaptive reconfiguration. Alternatively, a recent report showed that TIGIT could contribute to NK cell licensing [74]. In this paradigm, the presence of TIGIT may be required during NK cell maturation to ensure the acquisition of optimal effector function.

Some limitations should be noted in our study. First, the limited sample availability from a well curated study of untreated and treated HIV^+^ women only allowed us to profile NK cells from a small sample size of only female subjects. To overcome this, functional experiments were done on both female and male subjects. Additionally, the functional experiments were carried on healthy, HIV-negative subjects. This is particularly important for the blocking assays as viremic HIV^+^ individuals have higher TIGIT expression and thus may be more responsive to TIGIT blockade. Unfortunately, we did not have sufficient samples from viremic HIV^+^ patients to perform functional studies. Finally, our CyTOF panel did not exhaustively cover all known markers of adaptive NK cells and expression of PLZF, DAB2 and EAT2 was not assessed. CXCR6 was included in our original panel, but was excluded from analyses due to poor antibody staining.

In summary, our work demonstrates that TIGIT is overexpressed on NK cells of untreated HIV^+^ women, but fully corrected by ART. We find that TIGIT does not directly participate in the NK cell response to HIV-infected cells, but rather that TIGIT marks a mature NK cell subpopulation with adaptive features that is enhanced in functional responses to virus-infected cells, tumors, and cytokine stimulation. These results need to be confirmed in independent cohorts that evaluate both the direct functional contributions of TIGIT and the role of ART in regulating TIGIT expression.

## ACKNOWLEDGEMENTS

We are grateful to the Beninese study participants. We are indebted to N. Geraldo, A. Gabin, C. Assogba and C. Agossa-Gbenafa for their clinical expertise, to M. Massinga-Loembe, G. Ahotin, L.Djossou, and E. Goma for their technical assistance and to G. Batona and other field workers who helped with recruitment of commercial sex workers.

---

**Figure S1.**
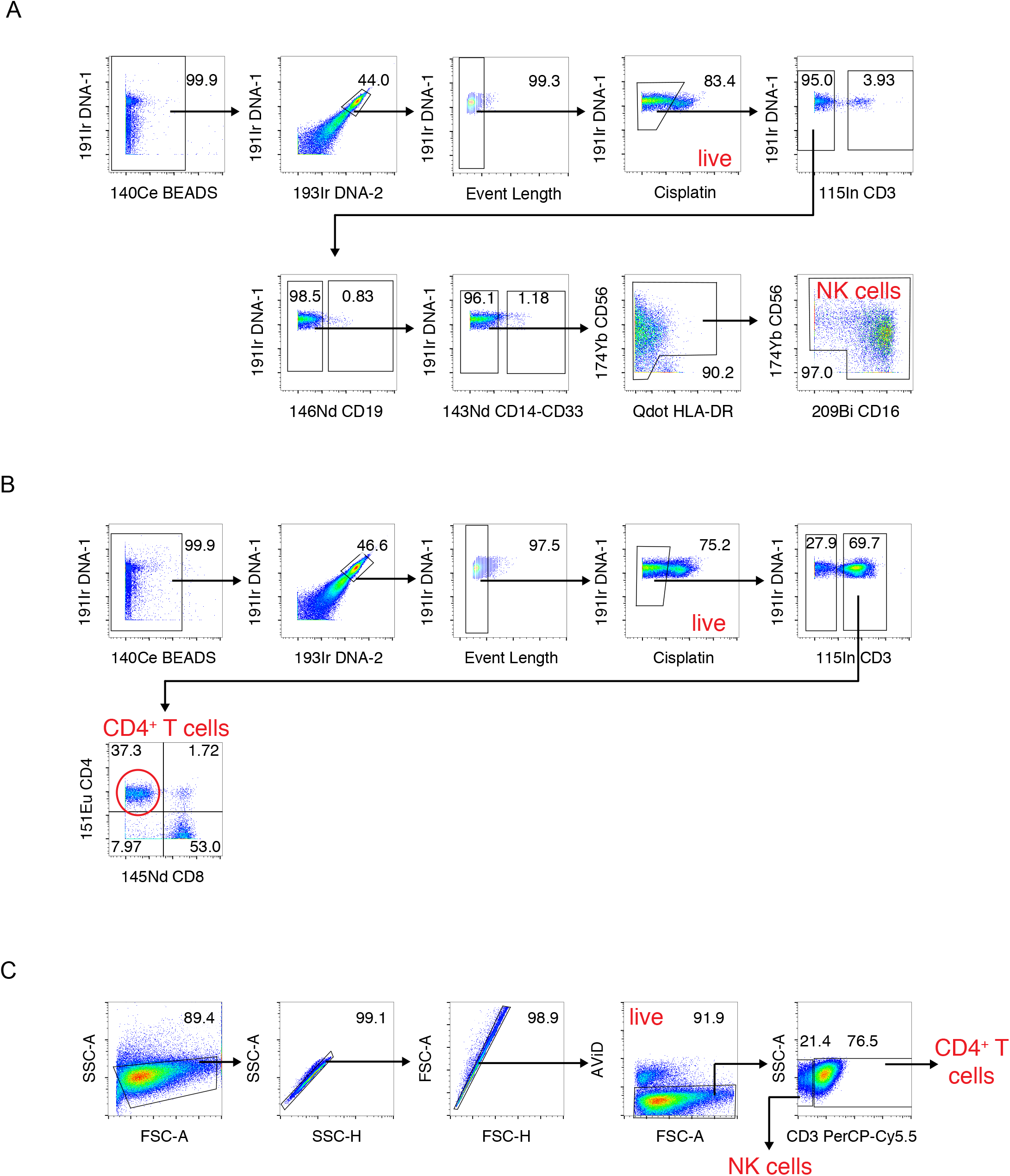
Gating Schemes. **(A)** Gating scheme used for negative gating of NK cells by CyTOF. NK cells were already purified with magnetic bead isolation prior to staining. Negative gating was performed to ensure further NK cell purity for downstream analyses. (B) Gating scheme used for analysis of CD4^+^ T cells by CyTOF. (C) Gating scheme used for analysis of NK cells and CD4^+^ T cells by flow cytometry.

**Figure S2.**
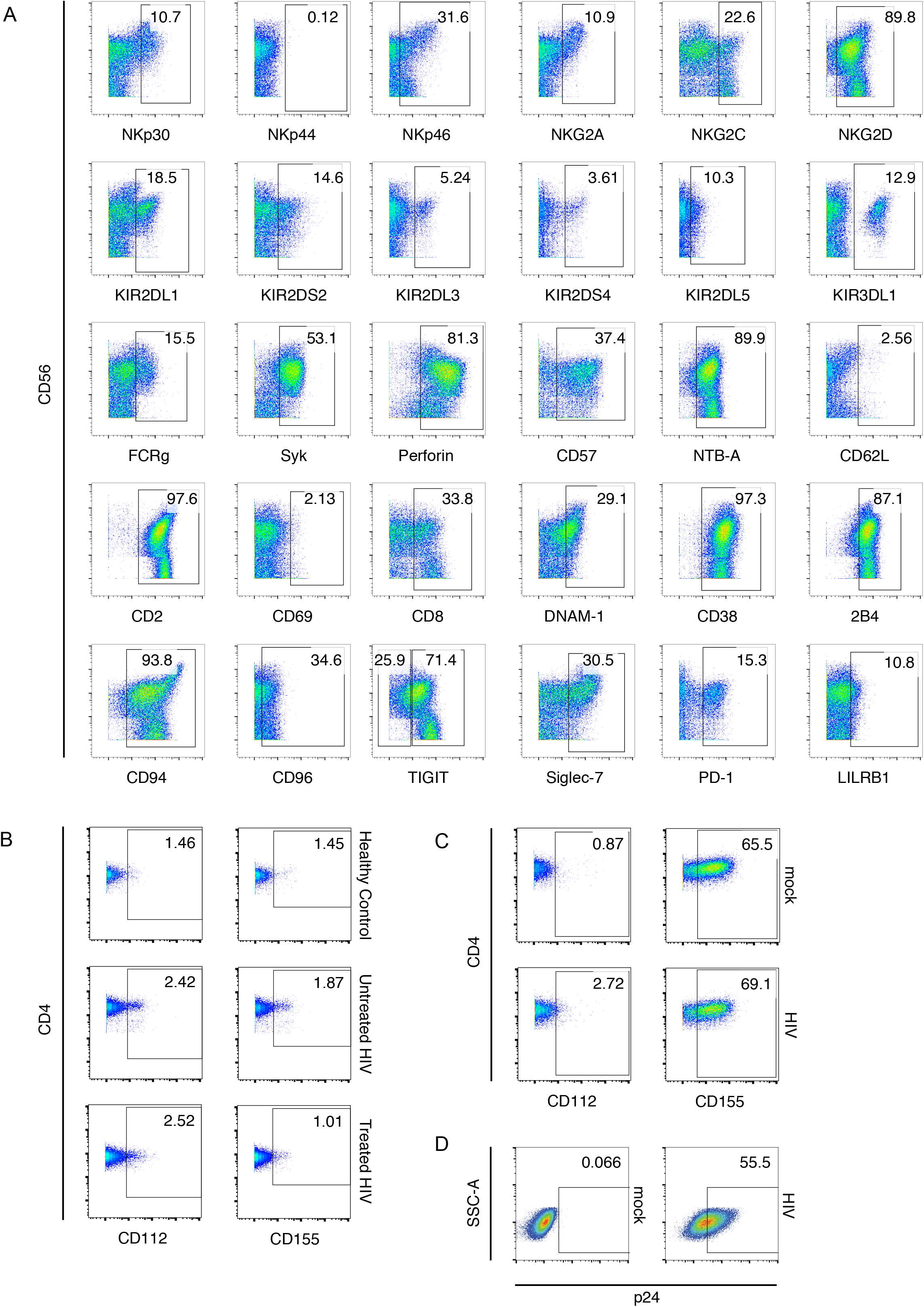
Gating strategy. **(A)** Representative CyTOF plots of the expression of NK cell markers from one of the treated HIV^+^ individuals. **(B)** Representative CyTOF plot of CD4^+^ T cell expression of CD112 and CD155 from one healthy woman, one untreated HIV^+^ woman and one treated HIV^+^ woman. **(C)** Representative CyTOF plots of CD112 and CD155 expression on mock-infected or HIV-infected CD4^+^ T cells. **(D)** Representative flow cytometry plot of p24 expression in mock-infected and HIV-infected CD4^+^ T cells. The median infection level for the blocking assay was 31.4% (range: 14.5-55.5%). The median infection level for the TIGIT^+^ vs TIGIT^-^ comparison was 38.0% (range: 14.6%-70.5%).

## SUPPLEMENTAL TABLES

**Table S1:**
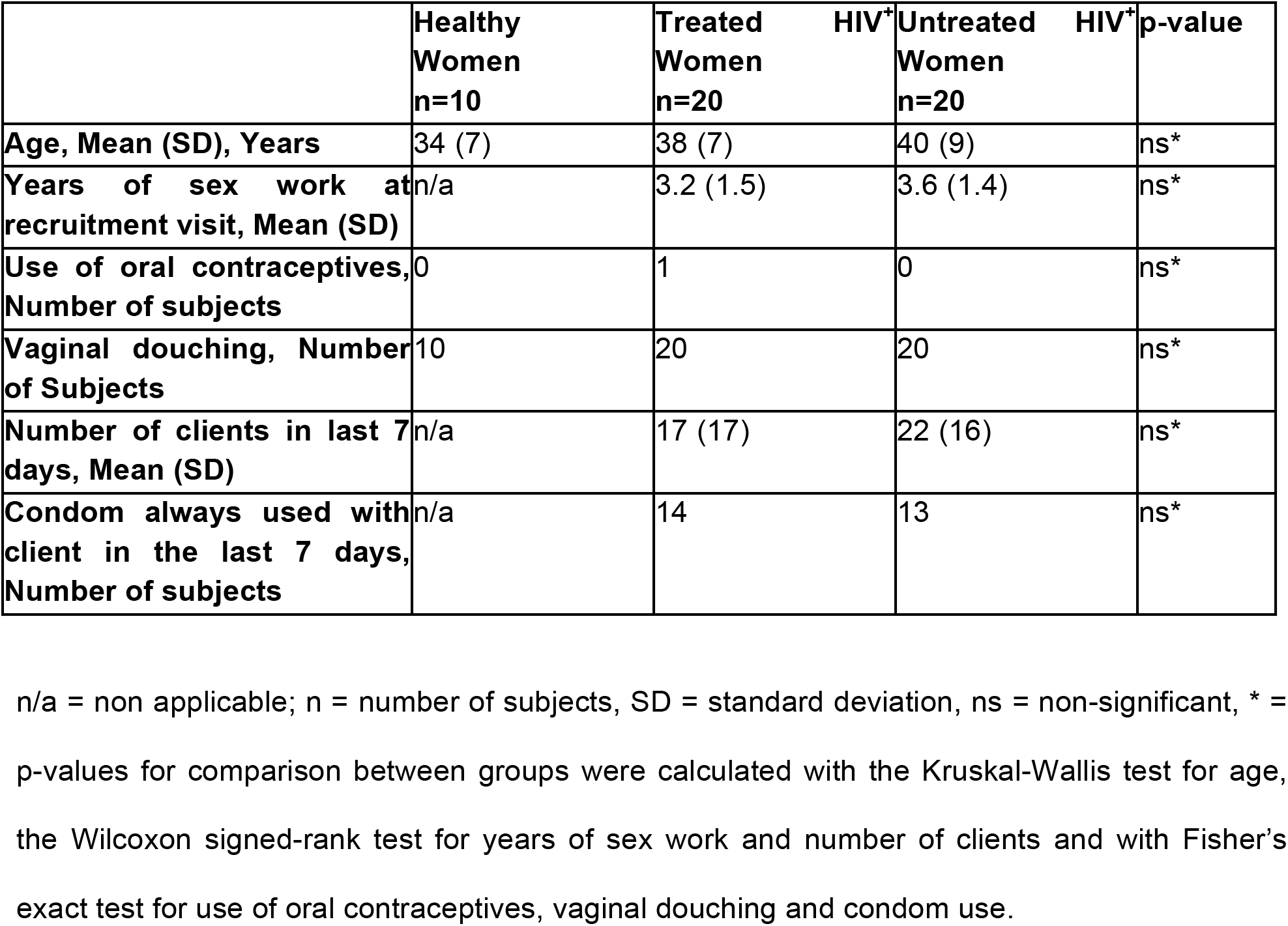
Study group demographics

**Table S2:** Ligand Panel

| **Isotope** | **Ligand Panel** | **Source** | **Clone** |
| --- | --- | --- | --- |
| 89Y | HLA-DR | Biolegend | L243 |
| Qdot | CD19 | Life technologies | SJ25-C1 |
| 115In | CD3 | Biolegend | UCHT1 |
| 139La | CD7 | Biolegend | 6B7 |
| 141Pr | CRACC | Biolegend | 162.1 |
| 142Nd | CD304 (BDCA-4) | Biolegend | I2C2 |
| 143Nd | PAN HLA | Biolegend | W6/32 |
| 144Nd | HLA-C | Millipore | DT9 |
| 145Nd | CD8 | Biolegend | SK1 |
| 146Nd | CD48 | Biolegend | BJ40 |
| 147Sm | CD303 (BDCA-2) | Fluidigm | 201A |
| 148Nd | ICAM1 (CD54) | Biolegend | HA58 |
| 149Sm | HLA-G | Biolegend | 87G |
| 150Nd | p24 | Abcam | ab9071 |
| 151Eu | CD4 | Biolegend | OKT4 |
| 152Sm | CD123 | Biolegend | 6H6 |
| 153Eu | CXCL13 | R&D Systems | 53610 |
| 154Sm | CCR2 | Biolegend | K036C2 |
| 155Gd | HLA-E | Biolegend | 3D12 |
| 156Gd | CD95 | Biolegend | DX2 |
| 157Gd | Nectin-1 (CD111) | Biolegend | R1.302 |
| 158Gd | MICA | R&D | 159227 |
| 159Tb | DR4 (CD261) | Biolegend | DJR1 |
| 159Tb | DR5 (CD262) | Biolegend | DJR2-2 |
| 160Gd | CD1c | Biolegend | L161 |
| 161Dy | ULBP-1 | R&D Systems | 170818 |
| 162Dy | CD11c | Fluidigm | Bu15 |
| 163Dy | ULBP-2-5-6 | R&D Systems | 165903 |
| 164Dy | Nectin-2 (CD112) | Biolegend | TX31 |
| 165Ho | PVR (CD155) | Biolegend | SKII.4 |
| 166Er | HLA-Bw4 | Miltenyi | REA274 |
| 167Er | NKp80 | Biolegend | 5D12 |
| 168Er | HLA-Bw6 | Miltenyi | REA143 |
| 169Tm | CD14 | Biolegend | M5E5 |
| 170Er | CD11b | Biolegend | ICRF44 |
| 171Yb | LFA-3 (CD58) | Biolegend | TS2/9 |
| 172Tb | CD33 | Biolegend | WM53 |
| 173Yb | CD141 | Fluidigm | 1A4 |
| 174Yb | CD56 | BD Pharmingen | NCAM16.2 |
| 175Lu | CD64 | Biolegend | 10.1 |
| 176Yb | MICB | R&D Systems | 236511 |
| 209Bi | CD16 | Fluidigm | 3G8 |

**Table S3:** NK Panel

| **Isotope** | **NK Panel** | **Source** | **Clone** |
| --- | --- | --- | --- |
| 89Y | CD57 | Biolegend | HCD57 |

| Qdot | HLA-DR | Life technologies | Tu36 |
| --- | --- | --- | --- |
| 115In | CD3 | Biolegend | UCHT |
| 141Pr | CD38 | Biolegend | HIT2 |
| 142Nd | CD69 | Biolegend | FN50 |
| 143Nd | CD33 | Biolegend | WM53 |
| 143Nd | CD14 | Biolegend | M5E5 |
| 144Nd | CD2 | Biolegend | RPA-2.10 |
| 145Nd | LILRB1 | Biolegend | GHI/75 |
| 146Nd | CD19 | Biolegend | HIB19 |
| 147Sm | CD8 | Biolegend | SK1 |
| 148Nd | FcRg | Millipore | Polyclonal |
| 149Sm | CD4 | Biolegend | SK3 |
| 150Nd | Syk | Biolegend | 4D10.2 |
| 151Eu | CD62L | Biolegend | DREG-56 |
| 152Sm | Ki-67 | Biolegend | Ki-67 |
| 153Eu | KIR2DS4 | R&D Systems | 179315 |
| 154Sm | KIR2DS2 | Abcam | Poly-clonal |
| 155Gd | NKp46 | Biolegend | 9E2 |
| 156Gd | NKG2D | Biolegend | 1D11 |
| 157Gd | TIGIT | R&D systems | 741182 |
| 158Gd | 2B4 | Biolegend | C1.7 |
| 159Tb | DNAM-1 | BD biosciences | DX11 |
| 160Gd | FAS-L | Biolegend | NOK-1 |
| 161Dy | NKp30 | Biolegend | P30-15 |
| 162Dy | Siglec-7 | Biolegend | S7.7 |
| 163Dy | NKG2C | R&D Systems | 134522 |
| 164Dy | NKp44 | Biolegend | P44-8 |
| 165Ho | CD96 | Biolegend | NK92.39 |
| 166Er | KIR2DL1 | R&D Systems | 143211 |
| 167Er | CD94 | Biolegend | DX22 |
| 168Er | CXCR6 | Biolegend | K041E5 |
| 169Tm | PD1 | Biolegend | EH12.2H7 |
| 170Er | KIR2DL5 | Miltenyi | UP-R1 |
| 171Yb | NKG2A | R&D Systems | 131411 |
| 172Tb | NTB-A | Biolegend | NT-7 |
| 173Yb | KIR3DL1 | BD biosciences | DX-9 |
| 174Yb | CD56 | BD biosciences | NCAM16.2 |
| 175Lu | KIR2DL3 | R&D systems | 180701 |
| 176Yb | Perforin | Abcam | B-D48 |
| 209Bi | CD16 | Fluidigm | 3G8 |

